# Validation of the Munich Actimetry Sleep Detection Algorithm for estimating sleep-wake patterns from activity recordings

**DOI:** 10.1101/2021.01.28.428574

**Authors:** Ann-Sophie Loock, Ameena Khan Sullivan, Catia Reis, Teresa Paiva, Neda Ghotbi, Luisa K. Pilz, Anna M. Biller, Carmen Molenda, Maria Vuori-Brodowski, Till Roenneberg, Eva C. Winnebeck

**Author notes:** Correspondence should be addressed to Eva Winnebeck and Till Roenneberg, Institute of Medical Psychology, Goethestraße 31, 80336 Munich. **Author contributions** (using CRediT-contributor taxonomy): Conceptualization: TR, ECW Data curation: AKS, NG, LKP, CM, AMB, MVB, ECW Formal analysis: ASL Investigation: CR (PSG), TP (PSG), AKS, NG, LKP, CM, AMB, MVB, ECW Software: TR (MASDA) Supervision: ECW Visualization: ASL, ECW Writing – original draft: ASL, ECW Writing – review & editing: AKS, CR, NG, LKP, AMB, ASL, TR, ECW. **Funding** ASL received a stipend by the Max-Weber-Programm (Studienstiftung); AMB from the Graduate School of Systemic Neurosciences Munich; CR from the Fundação para a Ciência e Tecnologia (FCT) PhD research grants (PDE/BDE/114584/2016); LKP a fellowship from the Coordenação de Aperfeiçoamento Pessoal de Nível Superior (CAPES, Finance Code 001); NG research funding from the FoeFoLe program at LMU (registration No. 37/2013). **Data availability** The data that support the findings of this study are available on request from the corresponding author. The data are not publicly available due to privacy or ethical restrictions.

## Abstract

Periods of sleep and wakefulness can be estimated from wrist-locomotor activity recordings via algorithms that identify periods of relative activity and inactivity. Here, we evaluated the performance of our Munich Actimetry Sleep Detection Algorithm (MASDA). MASDA uses a moving 24-hour-threshold and correlation procedure estimating relatively consolidated periods of sleep and wake.

MASDA was validated against sleep logs and polysomnography. Sleep-log validation was performed on 2 field samples collected over 54 and 34 days (median) in 34 adolescents and 28 young adults. Polysomnographic validation was performed on a clinical sample of 23 individuals undergoing 1 night of polysomnography. Epoch-by-epoch analyses were conducted and comparisons of sleep measures via Bland-Altman plots and correlations.

Compared with sleep logs, MASDA classified sleep with a median sensitivity of 80% (*IQR* = 75-86%) and specificity of 91% (87-92%). Mean onset and offset times were highly correlated (r = 0.86-0.91). Compared with polysomnography, MASDA reached a median sensitivity of 92% (85-100%), but low specificity of 33% (10-98%), owing to the low frequency of wake episodes in the nighttime polysomnographic recordings. MASDA overestimated sleep onset (~21 min) and underestimated wake after sleep onset (~26 min), while not performing systematically different from polysomnography in other sleep parameters.

These results demonstrate the validity of MASDA to faithfully estimate sleep-wake patterns in field studies. With its good performance across day- and nighttime, it enables analyses of sleep-wake patterns in long recordings performed to assess circadian and sleep regularity and is therefore an excellent objective alternative to sleep logs in field settings.

## Introduction

Sleep is characterized by reduced or absent consciousness, perceptual disengagement, immobility, and a characteristic sleep posture (Grandner & Rosenberger, 2019). The current gold standard to objectively detect and monitor sleep in humans is polysomnography (PSG), detecting characteristic EEG-patterns, muscle tone, and eye movements. Despite its usefulness in many clinical and research settings and its rich detail, PSG recordings and analyses are laborious and expensive, and thus rarely performed over several nights on the same person and consequently not well-suited to studying habitual sleep-wake patterns (Grandner & Rosenberger, 2019; Van de Water, Holmes, & Hurley, 2011).

In turn, actigraphy or actimetry (as we prefer to call it) monitors states of immobility through the detection of movements by wrist-worn devices (Conley et al., 2019; Dick et al., 2010; Grandner & Rosenberger, 2019; Marino et al., 2013; Roenneberg et al., 2015; Toon et al., 2016). Pioneering work in the 1970s and 1980s has made long, continuous recordings possible and demonstrated that sleep times and duration can be estimated from such records by identifying periods of relative immobility (Borbély, 1986; Kripke, Mullaney, Messin, & Wyborney, 1978). Thus, sleep-wake patterns can be captured over days, months or even years (Borbély, Rusterholz, & Achermann, 2017) and analyzed for sleep regularity, weekly or even seasonal patterns, effects of interventions, or sleep-wake rhythm disturbances resulting from circadian rhythm disorders (Ancoli-Israel et al., 2003; Dick et al., 2010; Kantermann, Juda, Merrow, & Roenneberg, 2007; Roenneberg et al., 2015; Sadeh, 2011; Smith et al., 2018). In addition to providing long-term records of sleep-wake patterns that are a prerequisite for circadian analyses, actimetry is less expensive and less sleep-disturbing than PSG and captures sleep where it normally occurs (Ancoli-Israel et al., 2003; Conley et al., 2019; Dick et al., 2010; Marino et al., 2013; Toon et al., 2016; Tryon, 2004). The easy handling and at-home applicability may also result in higher participation rates than in PSG studies (Marino et al., 2013). Actimetry even poses an advantage over sleep logs, which can be cumbersome to fill in over a long period and require literacy, but have often been the only practical alternative to investigate the long-term structure of sleep-wake rhythms (Girschik, Fritschi, Heyworth, & Waters, 2012).

The performance of actimetry in comparison to sleep logs and PSG in sleep-wake detection depends on the sleep-wake scoring algorithm, the recording device, the study population, and, of course, the question at hand. Generally, validations against PSG indicate adequate estimation of time and duration of sleep episodes as long as individuals do not have severe sleep fragmentation or severe sleep disorders (Ancoli-Israel et al., 2003; Sadeh, 2011). In such studies, actimetry was repeatedly shown to be highly sensitive in detecting sleep (sensitivity) but quite insensitive in detecting wake (specificity), thus indicating a tendency for movement-based sleep-wake scoring to overestimate sleep and underestimate wake (Conley et al., 2019; Grandner & Rosenberger, 2019; Marino et al., 2013; Sadeh, 2011; Van de Water et al., 2011). Importantly, these studies were mainly based on nighttime recordings, where sleep (not wake) is the most abundant and probable state. Therefore, the underestimation of wake reflects the underestimation of wake close to sleep onset and of wake-disruptions of sleep, i.e. wake after sleep onset (WASO), when people may remain relatively immobile in their beds – and not necessarily the underestimation of wakefulness during the day when people tend to move more. Accordingly, in validations against sleep logs that included wakefulness during the day, actimetric sleep-wake detection performed well in both sleep and wake detection and thus allowed monitoring changes in sleep patterns over time (Iwasaki et al., 2010; Lockley, Skene, & Arendt, 1999; Santisteban, Brown, & Gruber, 2018).

Here, we present the validation of our sleep-wake scoring algorithm called Munich Actimetry Sleep Detection Algorithm (MASDA) (sometimes referred to as bin-sleep method previously; Roenneberg et al., 2015). Like the many other algorithms introduced since the first validated algorithm by Webster, Kripke, Messin, Mullaney, and Wyborney (1982), it weighs the movement values within an epoch of interest against previous and subsequent epochs (Grandner & Rosenberger, 2019). However, MASDA was specifically designed from a circadian perspective to prioritize the detection of general sleep-wake patterns over the detection of sleep-wake in each individual short epoch. Hence it operates on a 10-min resolution of movement counts (10-min analysis epochs), which we commonly use in circadian actimetry analyses (Roenneberg et al., 2015), employs a 24-h-moving threshold for primary sleep-wake detection and yields relatively consolidated sleep episodes via a secondary, dedicated correlation procedure. This was intended to enable easier analyses and better pattern recognition on a 24-h-scale and long-range recordings than the more common short-epoch-by-epoch approach.

In this study, we validated the sleep-wake scoring results of MASDA against sleep-log entries from two samples collected over multiple weeks in the field as well as against PSG in single laboratory nights in a clinical sample.

## Methods

### Sleep log sample

#### Participants

For the validation against sleep logs, we used two samples from previous studies, an adolescent sample and one young adult sample. The adolescent sample was collected over 9 weeks in 45 German high school students (mainly Caucasians), of which 34 participants (22 females, *M* = 16.7 y, *SD* = 1.2, range = 14-19) provided high-quality data in both their sleep logs and actimetry records (median of 54 days) and were used for further analyses (Winnebeck et al., 2020). The young adult sample was collected over 4-6 weeks in 30 German participants (mainly Caucasians), of which 28 (13 females, *M* = 22.8 y, *SD* = 3.6, range = 19-33) provided complete data across both methods (median of 34 days) and made up the final adult sample (Ghotbi et al., 2020). Approval for both studies was obtained by the Ethics Committee of the LMU Medical Faculty (517-15, 774-16), and all participants (and their guardians if applicable) provided informed consent.

#### Actimetry

Activity was recorded with wrist-worn devices (Daqtometer 2.4, Daqtix) that were worn continuously on the wrist of the dominant or non-dominant hand (participants’ choice). This choice was possible as we did not aim to estimate general physical activity for metabolic monitoring but to estimate activity patterns. These dual-axis accelerometers were set to sample static and dynamic acceleration at 1Hz. Activity counts (the sum of the linear differences of subsequent readings for each axis) were stored by the devices at 30-sec intervals as the mean of all counts in this interval.

#### Sleep log

The sleep logs for both samples were based on the μMCTQ (Ghotbi et al., 2020), a short version of the Munich ChronoType Questionnaire. Instead of asking participants to record their average sleep times for the last weeks separately for work and work-free days, as this questionnaire normally does, the μMCTQ was applied daily via an online platform (limesurvey.org) to record sleep onset and offset of the previous night.

### PSG sample

#### Participants

For the validation against laboratory PSG, a dedicated dataset was recorded at the CENC Sleep Medicine Center, Lisbon, Portugal. The original sample consisted of 50 participants, however, because of software and signal synchronization issues (see below), records from only 23 of these participants (9 females, *M =* 40.1 y, *SD* = 13.7 y, range = 21-80 y) were used for the analysis. Of these, 11 subjects were diagnosed based on the PSG recording as without any clinical sleep pathology (healthy), 4 with insomnia, 1 with parasomnia (REM Behavior Disorder), 4 with Circadian Rhythm Sleep-Wake Disorder (2 Delayed Sleep-Wake Phase Disorder and 2 Shift Work Disorder), and 3 with Sleep Related Breathing Disorder (Obstructive Sleep Apnea). The study was approved by the Lisbon Medical School Ethics Committee and all participants gave their written consent.

#### Actimetry

Participants wore wrist actimeters ≥24 hours before and after the laboratory PSG night for a total of 14 days. For the majority of participants, ActTrust devices (Condor Instruments) were used, for 2 participants the Actiwatch 2 (Phillips Respironics). Sensitivity analyses without data from Actiwatch-2-participants yielded equivalent results indicating that device differences did not drive results. For both devices, activity was sampled every second and stored in 30-second bins. To evaluate the agreement between PSG and actimetry, temporal synchronization between the two methods was established via a synchronized event marker that was placed in both recordings.

#### Polysomnography

Overnight polysomnography was performed with the Nicolet System (Viasys Healthcare) or the Domino Somnoscreen Plus (Somnomedics). The recorded parameters included: electroencephalography (F3-M2, F4-M1, C3-M2, C4-M1, O1-M2, O2-M1); left and right electrooculogram; submental electromyogram; bilateral tibial electromyogram; electrocardiogram; oronasal airflow with 3-pronged thermistors; nasal pressure with a pressure transducer; rib cage and abdominal wall motion via respiratory impedance plethysmography; arterial oxygen saturation with pulse waveform; and digital video and audio. The recording was scored from “lights off” to “lights on,” with lights off scheduled as close as possible to participant’s normal sleep schedule, aiming for a sleep period of eight hours. In the case of an interfering work schedule or a sleep disorder that prevented participants from staying asleep (e.g. insomnia), a sleep period of fewer than eight hours was tolerated. The recordings were manually scored in 30-sec epochs by trained sleep technicians according to the American Academy of Sleep Medicine specifications (AASM version 2.3, 2017).

To match the 10-min resolution underlying MASDA (see below), the PSG scoring at 30-sec resolution was aggregated into 10-minute intervals via the mode, i.e. the most prevalent sleep stage over 20 consecutive 30-sec epochs was assigned to each 10-min interval. These were subsequently converted to a binary categorization (0 = wake, 1 = sleep). The median length of the series was 47.0 10-min intervals (*IQR* = 42.5-51.0), i.e. 7.8 h. Of this, participants spent 41.0 (37.5-44.5) intervals asleep and 5.0 (2-8.5) intervals awake.

### The Munich Actimetry Sleep Detection Algorithm (MASDA)

MASDA (Roenneberg et al., 2015), formerly also referred to as bin sleep method, is a two-step procedure for binary sleep-wake scoring from activity counts, heuristically designed to yield relatively consolidated stretches of sleep or wake. The first step of sleep detection is a threshold procedure, in which all epochs (usually 10-min long) with activity counts below a given percentage of the 24-h centered moving average are classified as putative sleep. The default percentage we use is 15% but it can be adapted from 10-25% for specific populations or individuals with particularly low or high activity during wake or sleep. The second step of MASDA is a “cleaning” procedure consisting of a duration filter and a correlation procedure. The filter reclassifies any sleep epoch not part of a stretch of at least 30-min as wake to avoid misclassification of short periods of inactivity. This is followed by a correlation procedure that joins adjacent stretches of sleep epochs based on a test series of sleep episodes of varying lengths. For more details on the correlation procedure please refer to Roenneberg et al. (2015). The algorithm was originally implemented in C++ in the software ChronoSapiens (© 2020 Chronsulting UG; Roenneberg et al., 2015), but has also been included in the Python package pyActigraphy by Hammad et al. (https://zenodo.org/record/3973012#.X_hCW14o8n0).

### Activity data processing

Actimetry data were analyzed via our in-house software ChronoSapiens (Version 10). All activity records were analyzed via our standard 10-min resolution (Roenneberg et al., 2015); the data were aggregated into intervals of 10 minutes via the arithmetic mean upon import into the program. Periods of non-wear were identified based on participants’ self-reports (actimetry-logs) as well as based on stretches of consecutive zeros exceeding 100 minutes and excluded from the analysis (i.e. set to NA). If these stretches occurred at the beginning of the inactive period on multiple days in the same individual, these were taken to be sleep with hardly any movement and not replaced with NA.

MASDA was performed with a 15% threshold (20% for the adolescent sample) and the setting to perform correlation series for four 10-min bins past the last r_max_ was 4.

Since MASDA incorporates information from the surrounding 24 hours via the 24-h-moving average, it can be influenced - to a certain extent - by stretches of missing data. To avoid systematic effects on the sleep-wake scoring, sleep bouts were excluded from the analysis if i) ≥1 hour of missing data was present within 3 hours or ii) ≥ 4 hours of missing data within 15 hours before or after the sleep bout. Accordingly, sleep bouts within the first or last 15 hours of the recording were also excluded.

### Method comparisons

#### Epoch-by-epoch agreement

Sensitivity, specificity, predictive values, and overall accuracy served as measures of agreement between MASDA and sleep logs/PSG. Sensitivity means the proportion of “true” sleep epochs (according to sleep log/PSG) that are also identified as sleep by MASDA. Specificity is defined as the proportion of “true” wake epochs (according to sleep log/PSG) that are also rated as wake by MASDA. While sensitivity and specificity relate the classification of MASDA to the ground truth (PSG/logs), the predictive values describe the probability that an obtained classification by MASDA is correct, taking the relative prevalence of sleep versus wake into account: the positive predictive value (PPV) quantifies accurate ratings of sleep, the negative predictive value (NPV) accurate ratings of wake. Overall accuracy is defined as the proportion of all sleep log/PSG bins that are correctly classified by MASDA.

#### Sleep parameter agreement

Summary sleep parameters calculated from the epoch time courses included sleep onset and sleep offset for both the sleep log and PSG sample, as well as sleep period time (SP), wake after sleep onset (WASO), total sleep time (TST), and sleep efficiency (SE) for the PSG sample. For the sleep-log sample, which encompassed multiple weeks of recordings, the average sleep onset and offset times per person over the entire recording were used. These averages were calculated after eliminating naps and fusing adjacent sleep bouts to obtain a daily onset and offset of the main sleep episode (see Winnebeck et al. (2020) for details). For the PSG sample, sleep onset was defined as the first bin scored as sleep after PSG recording started, sleep offset as the last bin scored as sleep before the PSG scoring ended. SP was defined as the elapsed time between sleep onset and sleep offset. WASO was calculated via the number of wake bins within a sleep episode. TST was defined as SP minus the amount of WASO. Sleep efficiency was the proportion of TST relative to time in bed (i.e., the length of the individual PSG recordings).

Correlation analysis between sleep parameters from PSG and MASDA was performed either via Pearson product moment correlations or via Spearman rank order correlations if parameters were non-normally distributed according to Shapiro-Wilk test. The alpha-level was set to 0.05. Additionally, Bland-Altman plots were created to visually examine the systematics of potential deviations between the sleep parameters derived from the two methods.

All analyses were conducted using R 3.5.1 and 4.0.2 (R Core Team, 2020) with special packages including psych (Revelle, 2020), tidyverse (Wickham et al., 2019), and data.table (Dowle & Srinivasan, 2020). Plots were generated using ggplot2 (Wickham, 2016) in R and matplotlib (Hunter, 2007) in Python 2.7.16 (Python Software Foundation, 2001-2019).

## Results

For our validation of the Munich Actimetry Sleep Detection Algorithm (MASDA), we made use of 3 different samples. Two samples with long continuous field recordings (medians of 54 and 34 days), one of adolescent students (n=34) and one of young adults (n=28), provided the basis for assessing MASDA against sleep-log records. A clinical sample with overnight PSG (n=23) including both patients with various sleep disorders as well as healthy sleepers was used to assess the algorithm against PSG. Validation included both assessment of epoch-by-epoch agreement as well as comparisons of standard summary parameters.

### Validation against sleep logs

#### Epoch-by-epoch agreement

For each individual of the adolescent and young adult samples, the sleep/wake classification for each 10-min epoch was compared between MASDA and the sleep logs. Over all participants, MASDA reached a median accuracy of 87% (*IQR* = 84-89%), sensitivity of 80% (75-86%), specificity of 91% (87-92%), positive predictive value of 80% (76-85%), and a median negative predictive value of 90% (88-92%) (Fig. 1). MASDA thus performed adequately in recognizing sleep: 80% of sleep-log-rated sleep epochs were correctly identified by MASDA (sensitivity), and 80% of algorithm-determined sleep epochs were also rated as sleep epochs in the logs (positive predictive value). In these long, continuous recordings, the algorithm performed even better in recognizing consolidated wake epochs: sleep-log-rated wake was identified as wake by MASDA in 91% of epochs (specificity), and 90% of epochs classified as wake by MASDA were sleep-log-rated wake as well (negative predictive value). The distribution of the epoch-by-epoch agreement metrics across participants is provided in Figure 1. Importantly, we identified no systematic differences between the two samples in any of the metrics (Table 1).

**Figure 1.**
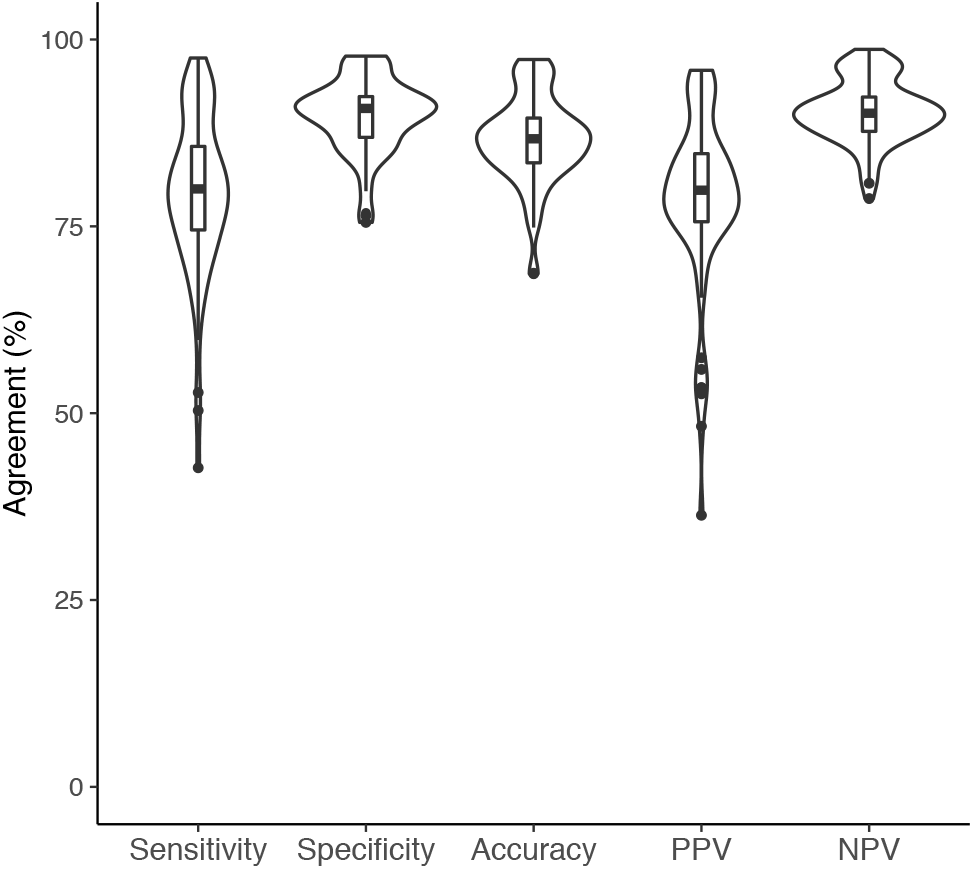
Epoch-by-epoch agreement between MASDA and sleep logs. Agreement of the sleep-wake scoring between both methods, with sleep log scoring as the ground truth, was assessed across multiple weeks in an adolescent and a young adult sample (total n=62). Results are displayed as a combination of violin and Tukey boxplots to illustrate data distribution. See Table 1 for the individual results of each sample. Abbreviations: PPV, positive predictive value; NPV, negative predictive value.

**Table 1.** Epoch-by-epoch agreement between MASDA and sleep logs – segregated by sample. Median agreement rates between the sleep-wake scorings by MASDA and sleep logs, with sleep-log scoring as the ground truth, are shown with their respective interquartile ranges (25^th^-75^th^ percentile) separately for the adolescent sample and the young adult sample. Results of Wilcoxon rank sum test comparing agreement metrics between both samples are also provided.

|  | **Adolescent sample**<br>Mean age = 16.7 y<br>n=34 | **Young adult sample**<br>Mean age = 22.8 y<br>n=28 | Sample comparison |
| --- | --- | --- | --- |
|  | *Median (IQR)* | *Median (IQR)* | *Wilcoxon test statistic* |
| **Overall accuracy** | 88% (84-90%) | 86% (83-89%) | W = 725, p = 0.29 |
| **Sensitivity** | 81% (76-90%) | 78% (72-83%) | W = 758, p = 0.15 |
| **Specificity** | 91% (87-93%) | 90% (88-92%) | W = 633, p = 0.98 |
| **PPV** | 80% (76-86%) | 79% (76-82%) | W = 687, p = 0.52 |
| **NPV** | 91% (89-92%) | 89% (86-92%) | W = 750, p = 0.18 |

#### Agreement in onset and offset measures

Agreement in onset and offset times between MASDA and sleep logs was assessed via correlations in the adolescent sample (Fig. 2). Correlating mean sleep onset times revealed a strong positive association (*R* = 0.92, *p* < 0.001). This association remained strong when differentiating between schooldays (*R* = 0.89, *p* < 0.001) and weekends (*R* = 0.91, *p* < 0.001). Mean sleep offset times also showed a strong relationship (*R* = 0.86, *p* < 0.001). Again, offsets were still significantly associated when differentiating between schooldays (*R* = 0.80, *p* < 0.001) and weekends (*R* = 0.86, *p* < 0.001), despite the systematic differences in wake-up times between these day types.

**Figure 2.**
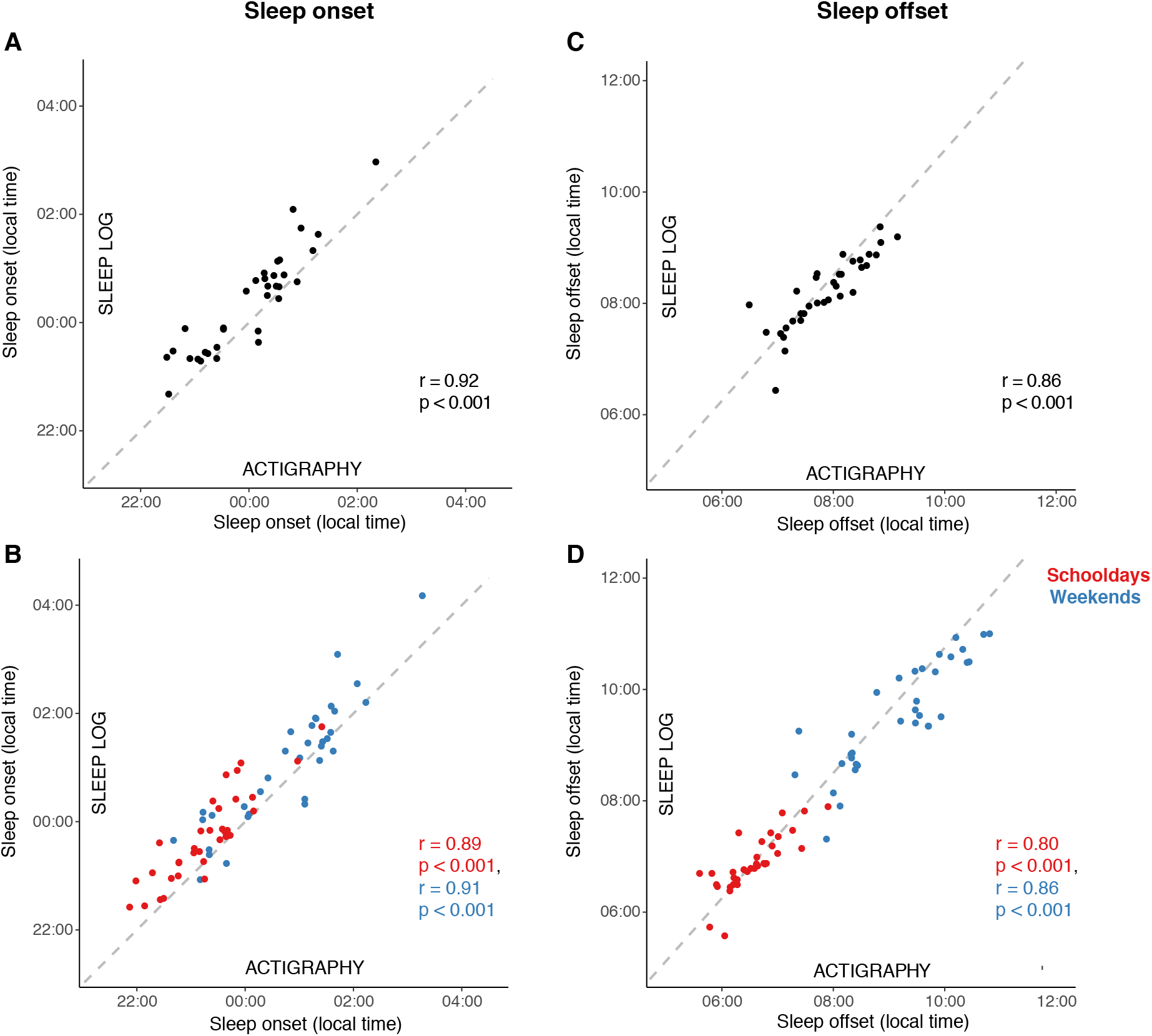
Relationship of sleep onset and offset from MASDA and sleep logs. Mean sleep onset times (A, B) and sleep offset times (C, D) of each participant from the adolescent sample (n=34) as determined via MASDA (x-axes) against those from sleep logs (y-axes). In panels A and C, means across all assessment days are compared, in B and D, the comparison is differentiated into means from schooldays (red) and weekends (green). Results of Pearson correlations are provided; dashed line represent a 1:1 relationship.

### Validation against PSG

#### Epoch-by-epoch agreement

Using the PSG sample, the sleep/wake classification of MASDA was compared to the PSG classification for each 10-min analysis epoch within each individual. Over all participants, MASDA reached a median accuracy of 83% (*IQR* = 78-92%), sensitivity of 92% (85-100%), specificity of 33% (10-98%), positive predictive value of 92% (87-99%), and negative predictive value of 37% (22-85%) (Fig. 3). Of note, specificity and negative predictive value both spanned the complete range from 0% to 100% (Fig. 3). In the PSG validation, where only nighttime sleep-wake states were assessed, MASDA thus performed best in detecting sleep: epochs considered sleep in the PSG scoring were identified as sleep by MASDA in 92% of cases (sensitivity), sleep epochs identified by MASDA were also PSG-identified sleep epochs in 92% of cases (positive predictive value). However, MASDA showed more difficulty in detecting wake: Only 33% of epochs that were considered wake by PSG were identified as wake by MASDA (specificity), and those that were classified as wake by MASDA were correct in only 37% of cases (negative predictive value). The lowest values for specificity and negative predictive value were in individuals with very few PSG-determined wake epochs. Here, misclassification by the algorithm weighed particularly strongly by definition.

**Figure 3.**
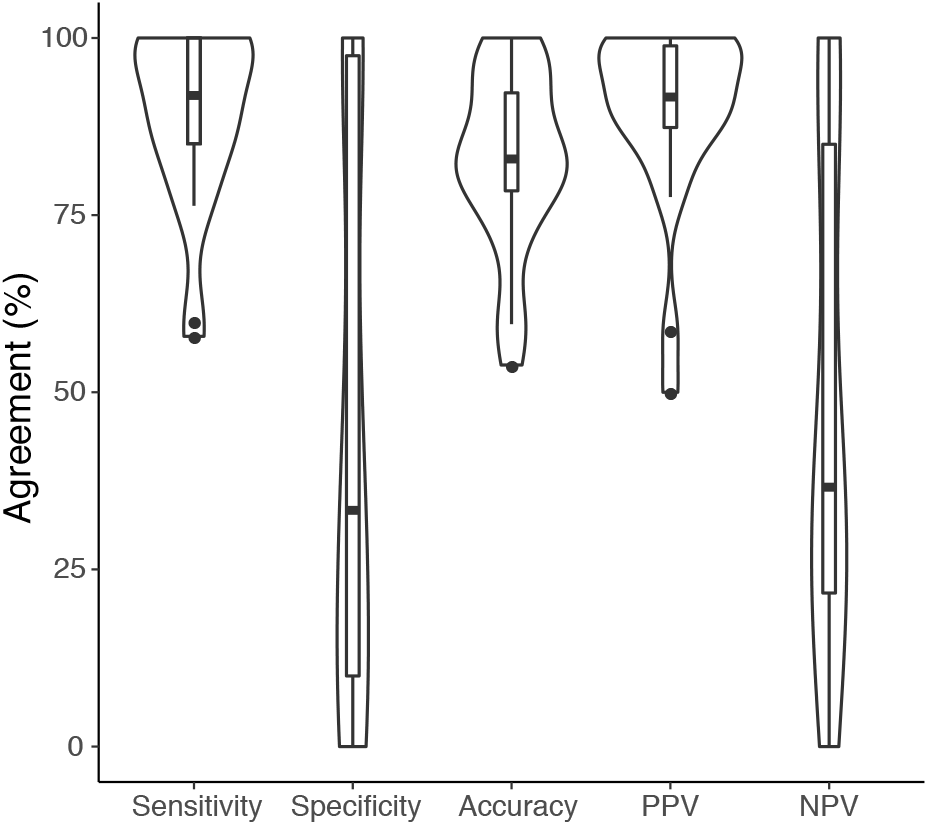
Epoch-by-epoch agreement between MASDA and polysomnography. Agreement of the sleep-wake scoring between both methods, with polysomnography as the ground truth, was assessed in single nocturnal recordings from a clinical sample (n=23). Results are displayed as a combination of violin and Tukey boxplots to illustrate data distribution. Abbreviations: PPV, positive predictive value; NPV, negative predictive value.

#### Agreement in sleep parameters

In addition to the epoch-by-epoch comparisons, we also analyzed agreement in common summary sleep parameters. Spearman correlation analyses between the parameters of both methods revealed a strong relationship for sleep onset (*rho* = 0.63, *p* = 0.01) and for sleep offset (*rho* = 0.76, *p* < 0.001). In contrast, sleep period duration, total sleep time, wake after sleep onset, and sleep efficiency, which more heavily depend on wake-detection during the sleep episode, showed no statistically significant relationship between the actimetry-determined and the PSG-determined values (Table 2).

**Table 2.**
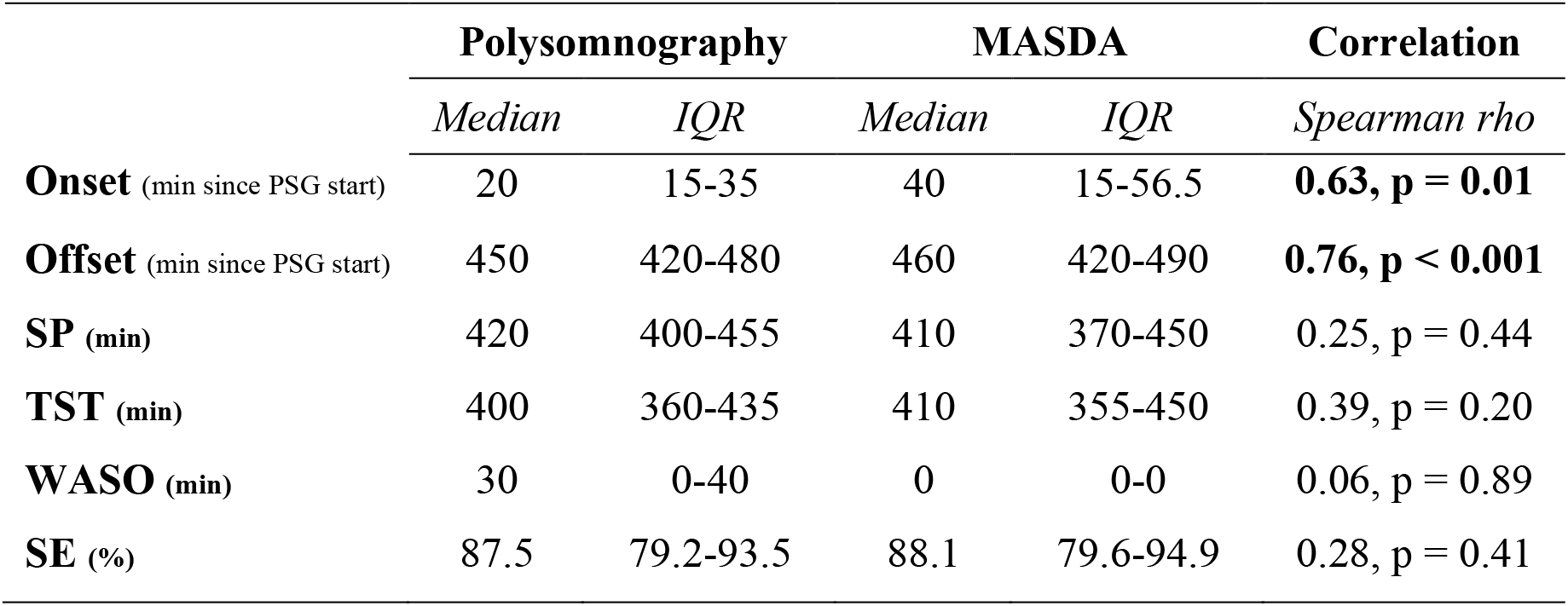
Summary sleep parameters from MASDA and polysomnography. Summary statistics of the summary sleep parameters from the clinical sample (n=23) are provided for both methods as well as the results of Spearman correlations between the two methods (p values are adjusted using the Benjamini-Hochberg correction). Abbreviations: SP, sleep period; TST, total sleep time; WASO, wake after sleep onset; SE, sleep efficiency.

Furthermore, Bland-Altman plots were used for visual inspection of potential systematic disagreements between actimetry-determined and PSG-determined summary measures (Figure 4). Bland-Altman analysis of sleep onset (Figure 4A) showed that onset times from MASDA were on average 21 minutes later than those from PSG. This pattern became more pronounced the later the sleep onset occurred (in relation to the start of the PSG recording). In contrast, sleep offset times for both methods (Figure 4B) were very similar, regardless of the relative timing of the sleep offset, with the bin sleep-detected offset deviating on average by less than 1 minute from PSG-determined values. In accordance with these later onset and similar offset times, the duration of the sleep period (from initial sleep onset until final sleep offset; Figure 4C) was on average 20 minutes shorter in MASDA. MASDA also underestimated wake after sleep onset (WASO; Figure 4D), as was expected from the lower specificity values obtained in the epoch-by-epoch analyses. WASO from MASDA was on average 26 minutes shorter than from PSG, notably showing outliers in both directions and no obvious relationships between WASO amount and method deviance. Lastly, both total sleep time (TST; Figure 4E) and sleep efficiency (SE; Figure 4F) deviated on average only marginally between the two methods. MASDA equally under- and overestimated TST and SE, while the deviance did not show any dependency on the amount of TST and SE.

**Figure 4.**
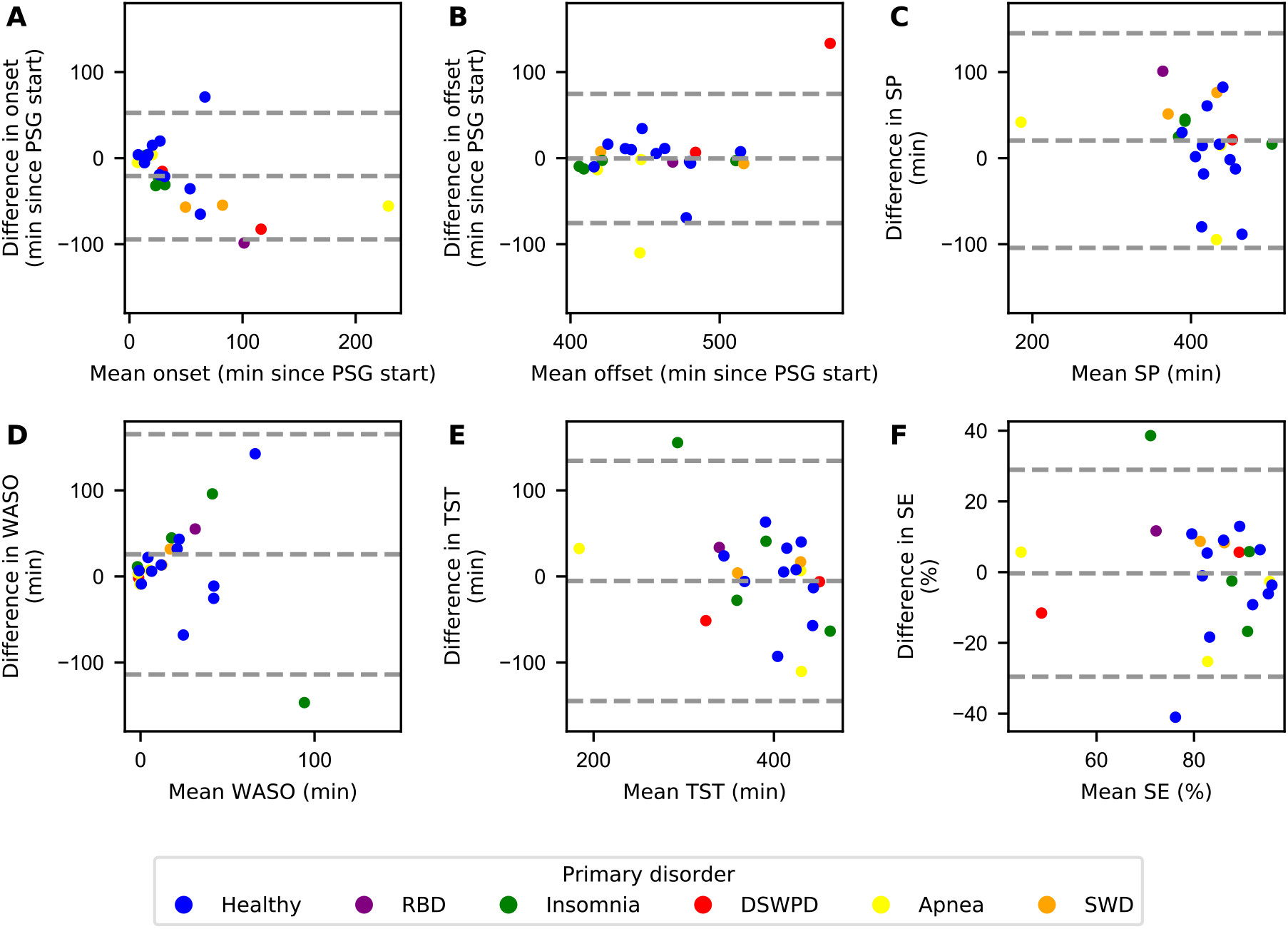
Bland-Altman analysis of summary sleep parameters from MASDA and polysomnography. Bland-Altman plots for (A) sleep onset, (B) sleep offset, (C) sleep period (SP), (D) wake after sleep onset (WASO), (E) total sleep time (TST), and (F) sleep efficiency (SE). In each panel, the mean of the sleep parameter between PSG and MASDA is plotted on the x-axis, the absolute difference in sleep parameters (PSG-MASDA) is plotted on the y-axis. The mean difference is denoted by the dashed line in the middle, the upper and lower boundaries of the 95% confidence interval by the upper and lower dashed lines. Colors indicate the primary sleep disorder diagnosed after the recording night as indicated in the legend. Abbreviations: RBD, REM Behavior Disorder; DWSPD, Delayed Sleep-Wake Phase Disorder; SWD, Shift Work Disorder.

Taken together, despite differences between the two methods, the Bland-Altman plots suggest that MASDA did not perform systematically worse than PSG in this one-night recording in most sleep parameters, except for sleep onset and WASO estimation. A few outliers in the difference between MASDA and PSG scores can be detected in most parameters, especially from one individual with delayed sleep-wake phase disorder (DSWPD). Nonetheless, sleep disorders do not seem to have systematically affected MASDA results as far as can be judged from this small sample (see color-coding in Figure 4).

## Discussion

In our comparing of the Munich Actimetry Sleep Detection Algorithm (MASDA) to sleep logs and PSG in three samples, we observed adequate rates of agreement throughout. In reference to sleep logs, MASDA performed well in detecting both consolidated sleep as well as consolidated wake states, and sleep onsets and offsets from MASDA and sleep logs correlated highly. In reference to a single night of PSG recordings, MASDA correctly identified sleep in most cases, yet showed a lower performance in detecting wake, i.e. the short, more instable states of wake right before and after sleep onset. MASDA deviated most from PSG in the assessment of sleep onset, WASO, and sleep period, while sleep offset, total sleep time and sleep efficiency were not systematically different between the two methods.

The good agreement of MASDA and sleep logs supports previous findings suggesting a reasonable validity between actimetry and sleep logs (Iwasaki et al., 2010; Santisteban et al., 2018; Usui et al., 1999). MASDA validation against PSG was also in line with other validation studies that have likewise noted an overestimation of sleep and underestimation of wake by actimetry in nocturnal recordings (e.g. Conley et al., 2019; Marino et al., 2013). Specifically, high sensitivity and overall accuracy rates have been reported in numerous validation studies (e.g. Marino et al., 2013; Van de Water et al., 2011). Poor specificity has frequently been reported as a problem of actimetry as well (Dick et al., 2010) and is thus only partly due to the algorithm’s design favoring consolidated stretches of sleep. Notably, we observed specificity and NPV rates ranging from extremely poor performance to perfect concordance with the PSG ratings.

Our results suggest that the comparison of actimetry to a night of PSG is not necessarily appropriate to evaluate the method’s 24-h-performance. The poor specificity rates obtained in the validation of various actimetry sleep-detection algorithms have often been brought up as a weakness of the method. However, as we show here, actimetry is not by definition worse at detecting wake states. In the sleep-log samples, which include both daytime and nighttime data across many days, our algorithm demonstrated good performance in both detecting consolidated sleep (i.e. good sensitivity/PPV), as well as consolidated wake periods (i.e. excellent specificity/NPV). The low specificity/NPV rates in the PSG sample likely result from the analysis of only about eight hours that are almost exclusively spent in bed, containing mainly sleep epochs, and very few wake epochs, while the wake epochs are marked by little activity. Indeed, if we assume the plausible scenario that all individuals from the PSG sample were continuously awake 3 hours prior to the start of the PSG recording and add these 3 hours of wake to the actimetry and PSG records, the rates for sensitivity and PPV remain the same, but median specificity increases drastically from 33% to 86% (*IQR* = 81-99%) and NPV from 37% to 86% (77-100%). Although we cannot be sure that all additional wake epochs would have been recognized as such by MASDA, this thought experiment illustrates the inherent bias towards low wake-detection if only nocturnal recordings are analyzed. Poor specificity values obtained under such conditions may accurately represent the method’s difficulty in identifying brief wake interruptions during sleep but not the method’s ability to identify longer wake episodes marked by more activity as occurring during the day.

The difficulty of correctly identifying wake interruptions during sleep was also evidenced by the underestimation of WASO by MASDA. In line with previous research stating that actimetry tends to overestimate sleep and underestimate wake during a sleep episode (e.g. Ancoli-Israel et al., 2003; Van de Water et al., 2011), this finding was particularly expected considering the 24-h-moving-average threshold employed by MASDA. The threshold heavily depends on daytime activity levels and thus trades the underestimation of short sleep interruptions for a high sensitivity for consolidated sleep-wake classification (Roenneberg et al., 2015). In line with Marino et al. (2013), we also noted a systematic increase in this underestimation, the longer the average WASO.

Likewise, the observed delay in sleep onset classification by MASDA in comparison to PSG is likely not random. Tryon (2004) introduced the idea of systematic differences between onset scorings since sleep onset has to be understood as a gradual change from wake to sleep. Actimetry typically marks the beginning of a sleep period by immobility (three 10-min bins of immobility under the threshold in our method), while PSG considers stereotypical changes in the electrical brain activity measured at the scalp, which can occur later (Marino et al., 2013; Tryon, 2004) or earlier.

Several limitations have to be put forward in interpreting our results. First, the PSG assessment was conducted in a laboratory environment, which can influence individual sleeping patterns. This setting also called for clearly defined in-bed intervals, which can limit the generalizability of the validation results (Grandner & Rosenberger, 2019), especially the high agreement in regard to sleep offset. Second, the validation was performed in rather homogeneous samples, so the results may not generalize to other populations, particularly people who move very little during the day (bed-ridden or elderly). The sleep-log validation was carried out in young, likely healthy, sleepers, while the PSG-validation was performed in a clinical sample where >50% of participants were diagnosed with sleep disorders. Unfortunately, the PSG-sample was diminished from 50 to 23 individuals due to software issues and hence we could not analyze effects of particular sleep disorders on the algorithm’s performance. Third, our analyses were performed on a resolution of 10 min, so each analysis epoch was only labelled with the most abundant state (sleep or wake) from the PSG 30-sec-epochs underlying it. This removed information about the relative proportion of sleep and wake within each analysis epoch, precluding the differentiation of performance between clear epochs and “swing epochs”. The 10-minute filtering that we generally apply to our human activity analyses has proven valuable when long-term, in-context-measures of daily sleep-wake behavior are investigated in contrast to the high-resolution architecture of single nights.

In conclusion, whilst PSG is undeniably richer in detail and more sensitive than the mere monitoring of body movements (Pollak, Tryon, Nagaraja, & Dzwonczyk, 2001), actimetry can be seen as an objective method of estimating sleep-wake patterns outside the laboratory (Meltzer, Montgomery-Downs, Insana, & Walsh, 2012), supporting large-scale, population-level sleep research. PSG and actimetry are suited for very different questions and settings, and thus are not in competition but should be seen as complementary to each other. By monitoring sleep longitudinally in natural settings, actimetry can help to detect sleep phase alterations, may assist in the diagnosis of circadian rhythm disorders (Ancoli-Israel et al., 2003; Smith et al., 2018) or the discovery of altered sleep patterns in individuals with sleep or neurobehavioral disorders (Sadeh, 2011). It can also provide objective data on treatment effects of non-pharmacologic and pharmacologic interventions (Ancoli-Israel et al., 2003; Brooks III, Friedman, Bliwise, & Yesavage, 1993; Roenneberg et al., 2015; Sadeh, 2011; Tryon, 2004). Especially when PSG measurements or sleep logs cannot be obtained, actimetry can contribute greatly to the understanding of individual sleep-wake patterns (Ancoli-Israel et al., 2003; Sadeh, 2011; Smith et al., 2018). We even use it to extract coarse patterns of sleep physiology from wrist-movements to assess sleep cycles in the field (Winnebeck, Fischer, Leise, & Roenneberg, 2018). Actimetry is thus a utile tool to measure sleep in diverse populations if conducted using validated algorithms (Smith et al., 2018). With our sleep detection algorithm, we hope to provide an additional useful and valid tool for studying sleep in the field. Given MASDA’s design to prioritize the detection of consolidated stretches of sleep over detecting frequent changes in sleep-wake state, MASDA lends itself not necessarily for detailed monitoring of WASO but all the more for circadian analyses striving to assess sleep timing and regularity.

## Notes

**Conflict of Interest:** TR, who developed MASDA, meanwhile uses MASDA in the context of the consulting work done in Chronsulting UG, of which he is the founder and CSO. All data were collected before TR started active work with Chronsulting. However, to avoid any potential construction of a conflict of interest, TR was not directly involved in this study’s data analysis or interpretation. He only provided general guidance and editorial input to the manuscript. In 2020, TR consulted for Estee Lauder Company; Vanda Pharmaceuticals; Jetlite; Salzgitter AG; NeuroCare; Vindex Company; KGK Science Ontario; and PricewaterhouseCoopers. All other authors declare that they have no potential conflicts of interest. During the conduit of this study, ASL received a stipend by the Max-Weber-Programm (Studienstiftung), NG received research funding from the FoeFoLe program at LMU, LKP a fellowship from the Coordenação de Aperfeiçoamento Pessoal de Nível Superior - Brasil (CAPES), AMB travel funds from the ESRS and the Graduate School of Systemic Neurosciences Munich, CR a stipend from the Fundação para a Ciência e Tecnologia (FCT) PhD research grants. During the conduit of this study but outside of the submitted work, ECW received travel and research support from the Friedrich-Baur-Stiftung, LMU Excellence, LMU Equal Opportunity, Gordon Research Conference and German Dalton Society.

### Competing Interest Statement

TR, who developed MASDA, meanwhile uses MASDA in the context of the consulting work done in Chronsulting UG, of which he is the founder and CSO. All data were collected before TR started active work with Chronsulting. However, to avoid any potential construciton of a conflict of interest, TR was not directly involved in this study's data analysis or interpretation. He only provided general guidance and editorial input to the manuscript. All other authors declare that they have no potential conflicts on interest.

